# Functional connectivity of the anterior and posterior hippocampus: differential effects of glucose in younger and older adults

**DOI:** 10.1101/482455

**Authors:** Riccarda Peters, David J. White, Brian R. Cornwell, Andrew Scholey

## Abstract

The hippocampus features structurally and functionally distinct anterior and posterior segments. Relatively few studies have examined how these change during aging or in response to pharmacological interventions. Alterations in hippocampal connectivity and changes in glucose regulation have each been associated with cognitive decline in aging. A distinct line of research suggests that administration of glucose can lead to a transient improvement in hippocampus-dependent memory.

Here we probe age, glucose and human cognition with a special emphasis on resting state functional connectivity (rsFC) of the hippocampus along its longitudinal axis to the rest of the brain. Using a randomized, placebo-controlled, double-blind, crossover design thirty-two healthy adults (16 young and 16 older) ingested a drink containing 25g glucose or placebo across two counterbalanced sessions. They then underwent resting state functional magnetic resonance imaging and cognitive testing. There was a clear dissociation in the effects of glucose by age. In older participants rsFC between posterior hippocampus (pHPC) and medial prefrontal cortex (mPFC) increased after glucose ingestion, whereas in younger participants connectivity decreased. Magnitude change in rsFC from pHPC to mPFC was correlated with individual glucose regulation and gains in performance on a spatial navigation task. Our results demonstrate that glucose administration can attenuate cognitive performance deficits in older adults with impaired glucose regulation and suggest that increases in pHPC-mPFC rsFC are beneficial for navigation task performance in older participants. This study is the first to demonstrate the selective modulation of pHPC connectivity in the acute setting.

---

Increasing the levels of available glucose, by the administration of a glucose drink, has been shown to improve cognitive performance in both younger and older adults in the minutes and hours following the drink ^1^. The effects of glucose have been reported to be comparable to those observed after administration of pharmaceutical cognitive enhancers ^2^. Converging evidence suggests a relationship between the effect of glucose on tasks which are predominantly related to hippocampal function such as episodic memory and spatial memory^1,2^.

The hippocampus is known to be important in the formation and recollection of memory ^3^and is a key brain hub for episodic memory, operating in the context of a large-scale network ^4^. Resting state functional connectivity (rsFC) analysis, including using functional magnetic resonance imaging (fMRI), has proved to be a powerful tool to help unravel the functional architecture of brain networks ^5^. One of the most common findings in studies of age-related rsFC is an association between advancing age and decreased functional connectivity within the default mode network ^6^ and overall reductions in functional connectivity between hippocampus and the rest of the brain ^7^.

It is increasingly recognized that there is a structural and functional dissociation between anterior and posterior segments of the hippocampus ^8^. This includes distinct patterns of resting state functional connectivity displayed by anterior (aHPC) and posterior parts (pHPC) of the hippocampus ^9^. The functional relevance of these networks and age-related changes therein are largely unknown. There is mixed evidence regarding the association between pHPC and aHPC connectivity and performance on cognitive tasks, and the functional differentiation of aHPC and pHPC is yet to be clearly defined. An episodic-spatial dichotomy of anterior and posterior hippocampal segments has been proposed, with pHPC being related to spatial memory functions and aHPC to episodic memory functions ^10^.

Much of the work on the influence of glucose on neurocognitive performance has focused on age-related effects ^11^. Senescence is accompanied by changes in glucose metabolism ^12^, specifically poorer glucose regulation. These changes in glucose regulation have been linked to age-related cognitive decline ^13,14^ and Alzheimer’s disease ^15^.

While cognitive domains implicated in the glucose facilitation effect have been argued to preferentially enhance hippocampus-dependent tasks, relatively limited research has directly explored underlying neurophysiological mechanisms in the human brain. Several event-related potential (ERP) studies support involvement of the hippocampus ^16-18^, further evidence stems from a study using fMRI ^19^.

To the best of our knowledge no study to date has considered the anterior-posterior division of the hippocampus in the study of the glucose facilitation effect. Furthermore, rsfMRI has not be directed to comparing the modulation of cognition and resting state connectivity in younger and older adults. The present study addresses these gaps in the literature. Here we describe the outcomes of a placebo-controlled, double-blind, crossover neuroimaging study investigating the relationship between age, glucose and human cognition with a special emphasis on the connectivity of the hippocampus along its anterior-posterior axis to the rest of the brain.

## Participants

A total of 32 healthy right-handed participants from Melbourne, Australia, were recruited for this randomised, double-blind, crossover trial. Half of this group (n=16, 8 women) consisted of younger subjects (mean age 25.8 ± 3.2 years, range 21-30) and the other half (n=16, 8 women) consisted of older subjects (mean age 68.6 ± 6.54, range 55-78). The participants were recruited via flyers, online advertising and from a database. All participants provided informed consent and received a small monetary compensation for their participation.

The study was approved by the Swinburne University Ethics committee and all procedures were performed in accordance with the principles of the 1974 Declaration of Helsinki.

Inclusion criteria included normal or corrected-to-normal vision and hearing, no major physical illness and had no history of neurological/psychiatric illness or head trauma. Further exclusion criteria were a diagnosis of diabetes mellitus, a history of hypersensitivity to glucose, heart disease or high blood pressure, smoking, substance abuse, intolerance to artificial sweeteners, pregnancy, claustrophobia, metal implants or any other contraindications to MRI.

Participants were excluded if they reported health conditions that would affect food metabolism including the following: food allergies, kidney disease, liver disease and/or gastrointestinal diseases (e.g. irritable bowel syndrome, coeliac disease, peptic ulcers). Subjects were also excluded if they were taking any medication, herbal extracts, vitamin supplements or illicit drugs which might reasonably be expected to interfere with blood glucose levels within four weeks prior to and during the study. In order to study associations with glucoregulation, the study included a range of fasting blood glucose levels. Fasting levels above 6 were presumed to reflect fasting compliance with compromised glucoregulation as long as they were noted consistently across visits. Where fasting levels above six were not noted across both visits, the subject was excluded from the analysis as this was taken as evidence of non-compliance with fasting.

During the testing session participants could be excluded if they scored lower than 25 on the Mini-Mental State exam (MMSE) ^20^.

All testing occurred at the Centre for Human Psychopharmacology at Swinburne University.

## Procedure

Participants attended the Centre for Human Psychopharmacology on three occasions a screening visit and two experimental sessions (with the experimental sessions balanced for treatment order). At the screening visit, informed consent was obtained, and eligibility was confirmed. Socio-demographic and morphometric data were collected (see **Table 1**). The session also served to familiarize participants with the cognitive tasks they would encounter during the study days.

**Table 1.** Demographics

Demographics
|  | Young | Older | *P*-value |
| --- | --- | --- | --- |
| **N (female)** | 16 (8) | 16 (8) | - |
| **Age (years)** | 25.0 ± 3.5 | 68.3 ± 6.4 | - |
| **BMI (m/kg<sup>2</sup>)** | 23 ± 4.5 | 25 ± 4.01 | 0.224 |
| **Education (years)** | 17 ± 2.0 | 15.4 ± 3.6 | 0.117 |
| **MMSE** | 29.8 ± 0.3 | 29.2 ± 1.0 | 0.025* |

*Note: for BMI, education and MMSE numbers represent: mean (standard deviation), significant results are marked \* at 0.05 level.*

The two experimental sessions started between 8.30am and 10am and were scheduled between two and 14 days, apart. Participants were asked to abstain from food and drink (except water) after 10pm before testing (to achieve a 12-h overnight fast). Fasting blood glucose levels were measured via capillary fingerprick using a Freestyle Optium Blood Glucose Sensor and Optium Blood Glucose Test Strips (Abbott Diabetes Care Ltd., Witney, UK) according to the manufacturer’s instructions. Following baseline glucose measurement, participants were asked to complete self-report questionnaires measuring mood and appetite. The self-report questionnaires were administered after each blood glucose measurement throughout the testing day and will be described elsewhere. They then received the treatment drink which consisted of 25 g glucose (Glucodin Pure Glucose Powder) mixed with 150 ml of water and 20 ml of sugar free coridal in the glucose condition, and two tablets (30mg) sodium saccharine (Hermesetas©) mixed with 150 mL of water and 20ml sugar free cordial in the placebo condition. It has previously been shown that the two drinks are indistinguishable in taste and mouthfeel ^21,22^. There is evidence that the glucose facilitation effect follows an inverted U-shape dose response curve in humans ^23^, and 25g has been reported as the optimal dose for memory enhancement ^2^.

Participants were randomly assigned to a treatment sequence that counterbalanced the order of treatments within age groups and gender. To ensure blinding, randomization and preparation of the drink were performed by a disinterested third party with no other involvement in the study. Blood glucose was measured again 20 min, 120 min and 150 min post-ingestion by a person who was not the experimenter to maintain blinding (unblinding occurred only after data analyses were completed).

MRI scanning commenced 30 min post-ingestion. This interval was selected to ensure that blood glucose levels would be maximally elevated during MRI scanning ^13^. Anatomical (T1) images were acquired first, followed by a seven-minute resting state scan.

## Cognitive Testing

Cognitive testing took place after MRI scanning (120 mins post-ingestion). Two cognitive tests focusing on different domains of memory were used, as follows.

### Working memory

Working memory performance was assessed using the mental arithmetic Serial Sevens task. It involves the serial subtraction of seven from a given number. Previous work has shown that performance on this to be enhanced by glucose administration^21^. Standard instructions were displayed on a computer monitor, informing the participant to count backwards in sevens from the given starting number, as quickly and accurately as possible, using the numeric keypad to enter each response. Performance was assessed using amount of correct subtractions within two minutes.

### Spatial learning and memory

Spatial learning performance was measured using a virtual analogue of the Morris Water Maze (vMWM) ^24^, the latter has been used extensively in rodents to study hippocampal-dependent spatial navigation.

Participants were instructed to try to find a platform in a virtual pool environment presented on a computer screen. The platform was visible on some trials and hidden (‘submerged’) on others.

Probe trials were administered in which the platform was removed unbeknownst to participants. In this case, participants started from a novel position in the pool in order to assess spatial memory performance beyond immediate training experience. The platform was then moved to a new location, and participants completed an additional set of hidden platform trials along with a second probe trial. The dependent measure was heading error (square-root transformed) or angular deviation from a straight path to the platform’s location on the probe trials.

For detailed task descriptions see **Supplementary material (S 1.1).**

## MRI acquisition

Participants underwent MRI scanning on both testing days, following the same protocol.

Functional magnetic resonance imaging (fMRI) data were acquired using a 3-Tesla Siemens Magnetom Trio scanner (Siemens, Erlangen, Germany) at Swinburne University of Technology, Melbourne, Australia with a 32-channel head coil. To minimize head movement comfortable padding was placed around the participants’ head and participants were instructed to lie as still as possible. Structural high resolution 3D T1-weighted magnetization prepared rapid acquisition gradient echo (MP-RAGE) anatomical images for anatomical reference were collected at the start of the scanning session (1mm isotropic MP-RAGE, TR= 1900 ms, TE = 2.52ms, flip angle= 9°). Following the anatomical image, the participant underwent 7 mins 15 sec of resting state fMRI (rsfMRI), during which they were instructed to keep their eyes open and look at a fixation cross. Functional data were obtained continuously with an interleaved multi band sequence (multiband acceleration factor= 6, bandwidth= 1860 Hz/Px, TR= 870ms, TE= 30ms, echo spacing= 0.69ms, flip angle= 55°, field of view= 192mm, voxel resolution= 2×2×2mm, slice orientation= transversal, number of slices= 66, 500 volumes per session). MRI scanning took 60 mins in total and also included two task related fMRI scans and a spectroscopy sequence that will be reported elsewhere.

## Resting state Analysis

Functional and structural images were processed using the CONN toolbox Version 17f ^25^; http://www.nitrc.org/projects/conn) for Statistical Parametric Mapping Software (SPM12; Wellcome Department of Imaging Science, Functional Imaging Laboratory, University College London) run under Matlab R2014a. Preprocessing steps were conducted using the default preprocessing pipeline for volume-based analysis (to MNI space) (realignment and unwarping, ART-based identification of outlier scans for scrubbing, simultaneous grey matter (GM), white matter (WM), and cerebrospinal fluid (CSF) segmentation and normalization into standard MNI space (Montreal Neurological Institute, Canada).

To remove confounding effects from the BOLD time series the anatomical CompCor strategy ^26^ was used as implemented in the CONN toolbox. Physiological and other spurious sources of noise were estimated and regressed out of the BOLD functional data in the denoising step (simultaneous option). Five principal components were extracted from both WM and CSF, as well as 12 motion regressors (six head motion parameters + six first-order temporal derivatives) derived from spatial motion correction were used as temporal covariates and removed from the BOLD functional data using linear regression. The resulting residual BOLD timeseries were band-pass filtered with a frequency window of 0.008 Hz-0.2Hz.

In addition to the above steps controlling for motion, an analysis of motion across groups and conditions was conducted. Framewise displacement (FD; maximum total and averages scan-to-scan) was calculated according to Power et al. ^27^ between-sessions and between-groups (see **Supplementary material 2.1**). As FD was higher for younger adults than older adults at both treatment sessions, and FD also differed between sessions in the younger group, all group level analyses included average framewise displacement as a covariate at the second level.

## Seed Based Connectivity analysis

To assess rsFC of the hippocampus, four seed regions were defined based on coordinates which have been used in previous research ^9^ to facilitate comparisons of results. Four ROIs were created in MarsBar toolbox (http;/marsbar.sourceforget.net/) and defined as spheres with a 5mm radius around the anterior and posterior part of the hippocampus on the left (anterior: x= −28, y= −12, z= −20; posterior: x= −28, y= −24, z= −12) and right side (anterior: x= 28, y= −12, z= −20; posterior: x= 32, y= −24, z= −12). The resting state BOLD signal timeseries of each hippocampal region of interest was extracted and correlated against voxels of the rest of the brain for each session of each subject, Fisher z transformation was applied.

## Statistical Analysis

Blood glucose level and cognitive outcomes were assessed by mixed design ANOVA, with treatment as a repeated measures factor and age as a between-subjects factor (IBM SPSS statistics, Version 24). Blood glucose levels had an additional repeated measures factor of assessment time within sessions. Glucose tolerance was assessed as incremental area under the curve (iAUC) using the trapezoidal rule ^28^ based on the four blood glucose measurements taken at baseline, 20 mins, 120mins and 150mins post glucose administration. Higher iAUC values reflect higher circulating glucose levels, indicative of poorer glucoregulatory ability.

Functional connectivity analyses used whole-brain voxel-wise mixed within subject (glucose, placebo) and between subject (younger, older) second-level models, using the partitioned variance approach implemented in CONN, in order to test for treatment x age-group interactions. All rsFC analyses used a cluster-extent FWE-corrected *p*-value < 0.05, obtained using non-parametric statistics with 5000 permutations at a cluster-defining threshold of *p* < 0.001.

To explore the relationship between changes in resting state functional connectivity and performance on cognitive measures, *post-hoc* Pearson’s correlation coefficient analyses were carried out using SPSS.

## Results

Data from two participants of the young group (one male, one female) was omitted from the analysis because they showed significantly higher fasting glucose levels at a single session compared to the other session.

## Blood glucose levels

Blood glucose levels throughout both testing sessions for each group are depicted in **Figure 1a**., which also presents the timing of the experimental measures. There were no between-group differences in baseline blood glucose levels between the younger and older group either at glucose (*T*(1, 28)= 0.59, *P*= 0.954) nor placebo visit (*T*(1, 28)= −0.878, *P*= 0.387). A 2 (Age: young/old) × 2 (Treatment: Glucose/Placebo) × 4 (Timepoint: 0, 20, 120, 150 min) ANOVA was conducted. The rmANOVA analysis of glucose levels omitted data from one young participant who had a missing post-dose blood glucose assessment at the placebo visit. There was a significant main effect of Treatment (*F*(1,27) = 25.18 *P* < 0.001, η2 = 0.485), a main effect of Timepoint (*F*(1.567, 27) = 59.43, *P* < 0.001, η2 = 0.688, Greenhouse-Geisser) and a significant Treatment × Timepoint interaction (*F*(1.843, 27) = 19.43, *P* < 0.001, η2 = 0.644, Greenhouse-Geisser). There was also a main effect of Age (*F*(1, 27)= 9.076, *P*= 0.006, η2 = 0.252), the older group showed a greater increase of blood glucose levels in response to glucose ingestion than the younger group (*T*(1, 28)= −2.99, *P*= 0.006).

**Figure 1.**
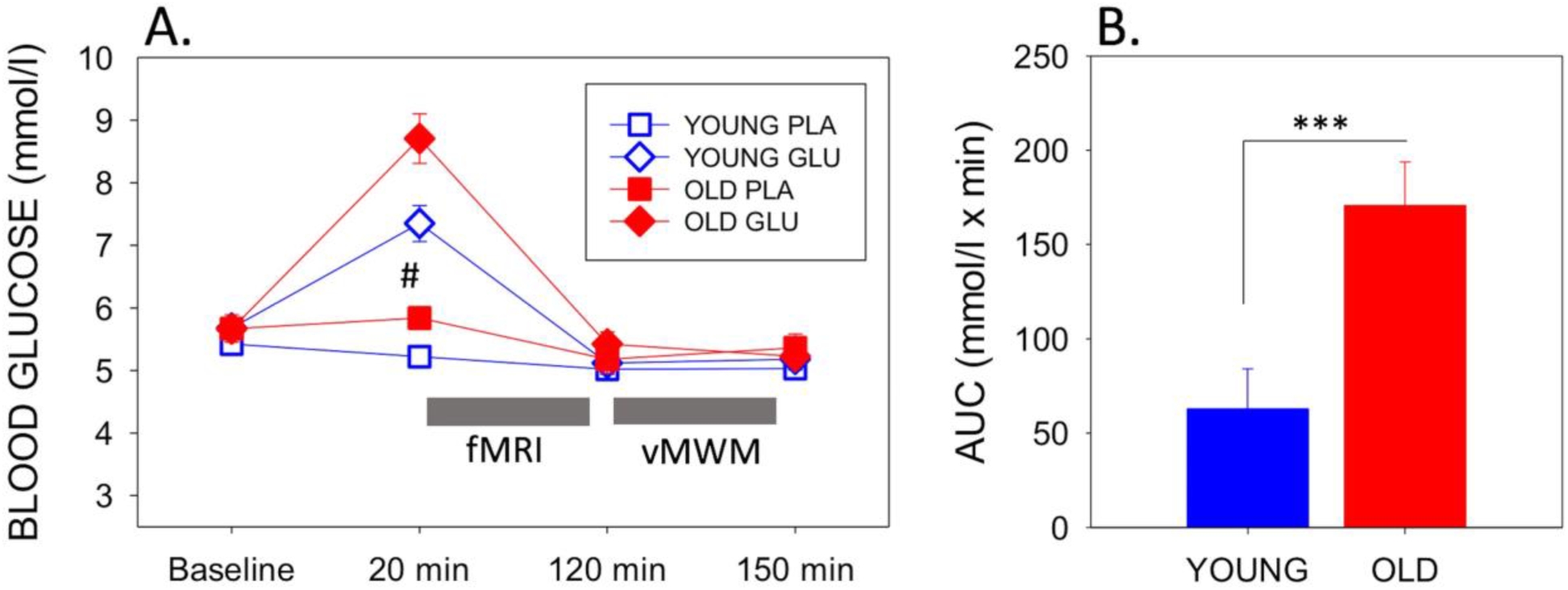
a) Mean (with SEM) blood glucose levels at baseline, pre-MRI (20mins post-dose), post-MRI (120mins post-dose), and end of testing (150mins post-dose) for each group and each visit. Circles depict younger adults, while squares depict older adults. Filled symbols represent measures taken on glucose visit, while open symbols represent measures at the placebo visit. # indicates significant difference between drink condition and significant difference between ages in both drink conditions (p-values see text). Timing of fMRI and virtual Morris Watermaze (vMWM) are indicated in relation to glucose measurements. b) Blood glucose incremental area under the curve as a measure of glucoregulatory efficiency, bars depict mean (with SEM). Older adults had significantly higher iAUC than younger adults (T(1, 28) = −3.403, P= 0.002), indicating poorer glucose regulation. *** P < 0.005.

There was also significant difference in blood glucose levels 20 mins post-ingestion at the placebo visit (*T*(1, 27)= − 3.72, *P*= 0.001), glucose levels decreased in the younger group. Using blood glucose iAUC at the glucose visit as a measure of glucoregulatory efficiency, older participants had significantly higher iAUC than younger participants (*T*(1, 28)= − 3.403, *P*= 0.002), indicative of poorer glucose regulation in the older sample (Figure **1b**).

## Resting State

A significant Treatment x Age-group interaction in rsFC was observed between left pHPC and a cluster within in the mPFC encompassing areas in anterior cingulate, paracingulate gyrus and superior frontal gyrus (Brodmann Area (BA) 32 and 8; MNI peak [+08 +26 +30] (**Table 2 & Figure 2a**).

**Table 2.**
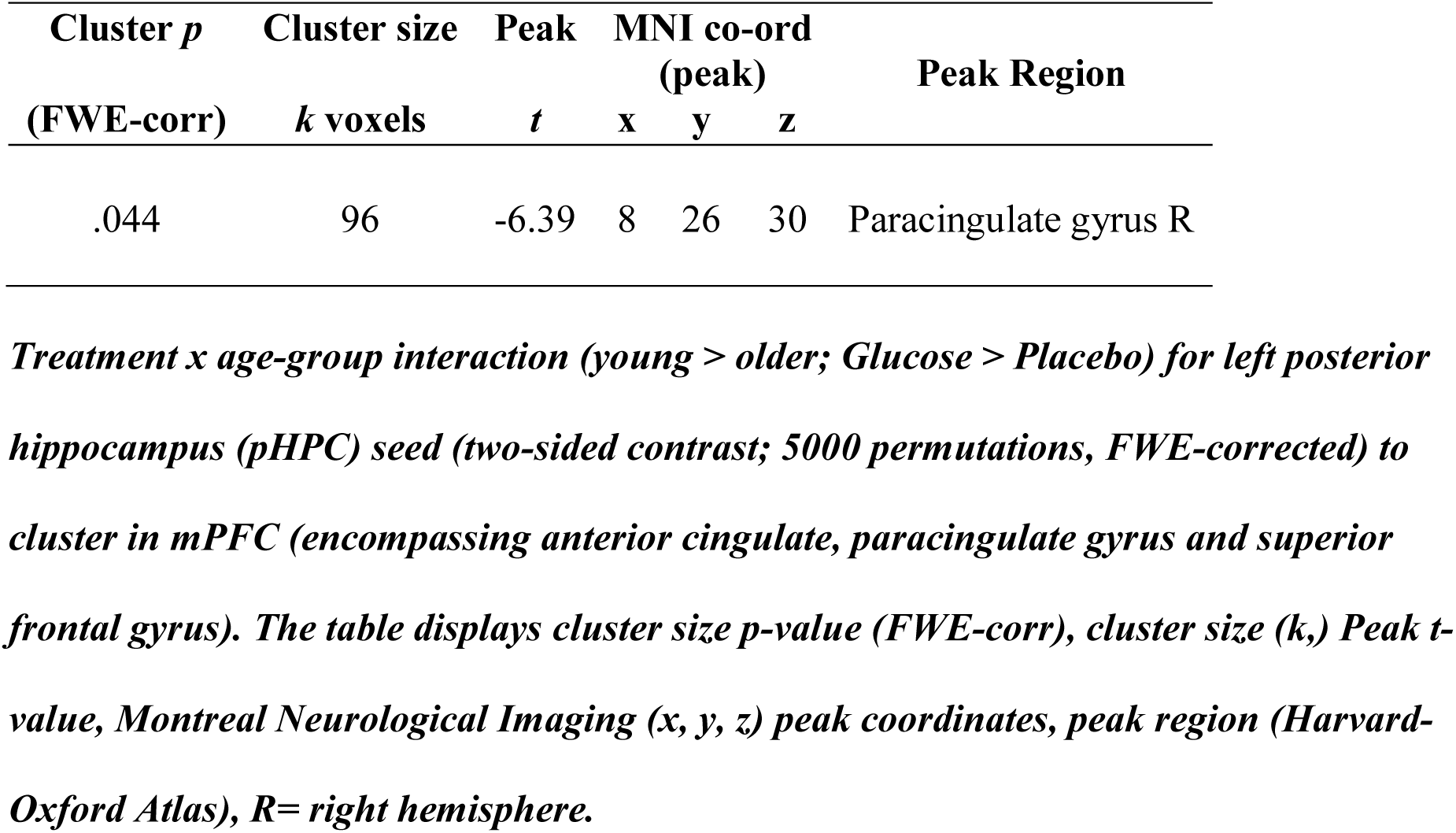
Whole brain voxel-wise rmANOVA of resting state functional connectivity with left posterior hippocampus

*Post hoc* t-tests revealed that glucose (compared to placebo) significantly increased left pHPC-mPFC connectivity in older participants, *T*(1,15)= 5.13, *P* <0.001), whereas the reverse was observed in young participants, *T*(1,13)= −3.6, *P*= 0.003 (**Figure 2b**).

**Figure 2.**
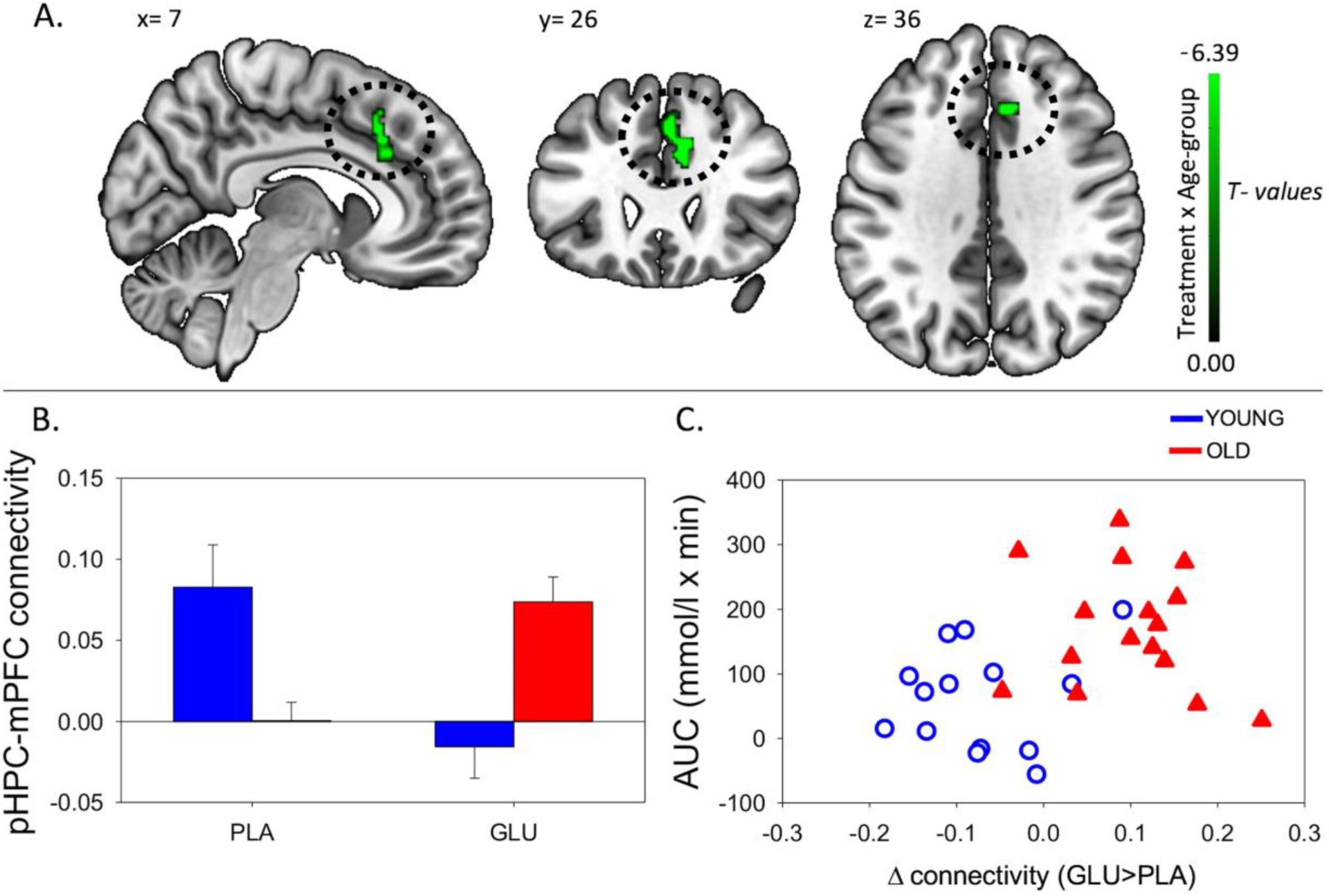
a) Resting-state functional connectivity brain map for left posterior hippocampus showing: Treatment x Age-group interactions exhibited from left pHPC to cluster in mPFC (encompassing anterior cingulate, paracingulate gyrus and superior frontal gyrus) (fisher z-transformed correlation values). b) Extracted connectivity strength from pHPC to mPFC for each group per session (error bars reflect SEM). c) Scatterplot of correlation between change in rsFC of pHPC and mPFC and glucose regulation as measured by iAUC (r = 0.39, P =0.04).

This analysis was repeated using individual pHPC volume as a covariate. The cluster in the same area remained significant although smaller in size (MNI peak [+04 +24 +44]; Voxels (k)= 58) (see **Supplementary Material 2.2**).

No significant differences were observed from any of the other seed ROIs (right pHPC, or left and right aHPC).

Further *post hoc* tests showed that younger participants exhibited higher rsFC between left pHPC and mPFC relative to older participants under placebo conditions, *T*(1,28)= 3.54, *P*= 0.001; and that older participants exhibited significantly greater left pHPC-mPFC connectivity relative to the younger group after glucose ingestion alone, *T*(1,28)= −2.75, *P*= 0.01 (**Figure 2b**).

To relate the present finding to individual glucose regulation, *post hoc* Pearson correlation using iAUC at the glucose treatment visit and rsFC connectivity were performed. Change in pHPC-mPFC rsFC was correlated with individual glucose regulation across the whole sample (r = 0.39, *P*=0.04) (**Figure 2c**).

## Task Performance

Data from two additional participants (one young, one old) was omitted from the analysis of the Morris Water Maze due to missing data. There was a significant Treatment x Age-group interaction for performance on the Morris Water Maze task (*F*(1, 26)= 8.64, *P*= 0.007) as depicted in **Figure 3a**. Posthoc pairwise comparisons revealed that while the older group showed significantly worse performance compared to the younger group under placebo, (*T*(1, 26) = −3.42, *P*= 0.02), there was no group difference after glucose administration, (*T*(1, 26)= 0.35, *P*= 0.73).

**Figure 3.**
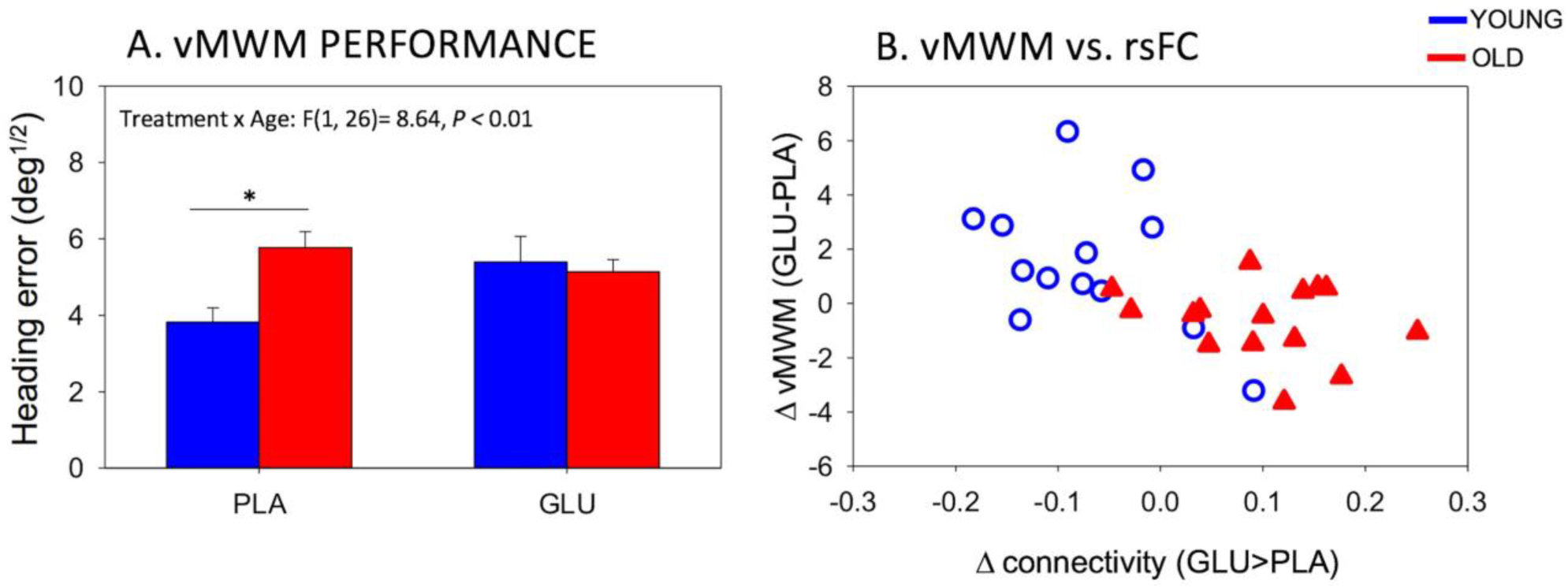
a) Behavioural performance on the virtual Morris water maze (vMWM) task for younger and older subject following placebo and glucose drink. Bars depict mean Heading error (deg^1/2^) (with SEM). Smaller values reflect better performance. b) Scatterplot of change in resting-state functional connectivity of left posterior hippocampus to mPFC and change in performance on the spatial navigation task (heading error) (r = −0.56, P = 0.002).

No Treatment by age-group interactions were found in the Serial Sevens (correct) (*F*(1, 28)= 1.38, *P*= 0.25).

We further probed the relationship between significant changes in left pHPC-mPFC rsFC and change in performance on the spatial navigation task.

Change in rsFC magnitude between left pHPC and mPFC was correlated with changes in spatial memory performance (r = −0.56, *P* = 0.002). The negative correlation reflects greater connectivity after glucose to be associated with smaller heading errors (averaged across the two probe trials) after glucose ingestion. Specifically, in older participants, glucose increases functional connectivity between left posterior hippocampus and mPFC and that the magnitude of this functional connectivity change correlates with the change in performance (**Figure 3b**).

## Discussion

The aim of the present study was to investigate age-related differences in changes in rsFC in response to a glucose load from the anterior and posterior segments of the hippocampus, comparing a group of younger and a group of older adults. Further, we wanted to investigate if these changes are linked to individual glucose regulation and how they relate to memory performance on two different memory tasks.

After the ingestion of a 25 g glucose drink, we observed changes in functional connectivity of the pHPC to a cluster in the mPFC. In older participants, glucose increased rsFC between pHPC and mPFC. Furthermore, the change in connectivity was related to glucoregulatory ability, participants with poorer glucose regulation, as indicated by greater circulating glucose following administration, benefited most from the glucose load. In younger participants we observed the opposite pattern of connectivity change. Furthermore, the change in magnitude of rsFC was correlated with gains in performance on a spatial memory task.

Here we demonstrate age-dependent, acute modulation of pHPC connectivity, suggesting that connectivity of the posterior segment of the hippocampus is differentially susceptible to an acute glucose load in older individuals.

This study is the first to compare the effects of glucose on rsFC and cognition in both younger and older individuals. There is a general consensus that glucose enhancement is more effective in older compared to younger adults ^29^. This effect has been partially attributed to an age-related decline in glucose regulation. Our findings indicate that pHPC-mPFC connectivity increases were more marked as glucoregulation worsened in older participants (see **Figure 2c**). Our results suggest that these connectivity changes may contribute to previously reported demonstrations that glucose more readily attenuates age-related cognitive performance decrements in elderly adults with impaired glucose regulation^30,31^.

The hippocampus has been hypothesised as a key structure in the glucose facilitation effect ^1^. Our results show the potential importance of considering subdivisions of the hippocampus along its anterior-posterior axis in neuropharmacological studies.

The hippocampus and surrounding temporal lobes have long been recognised to be an important node for the processing of spatial information (for a review, see ^32^). The results of the present study support the role of the posterior segment of the hippocampus in spatial memory performance. This relationship has recently been demonstrated in a rsfMRI study by Persson et al. ^10^, who predicted spatial memory performance from pHPC, but not aHPC, rsFC. Our results are also consistent with reports of positive associations between posterior hippocampal activation and virtual navigation performance ^33-35^.

Growing evidence points to the importance of the medial frontal cortex (mPFC), including the anterior cingulate cortex (ACC), in spatial memory. Interactions between mPFC and hippocampus have been proposed to be critical to successful encoding and retrieval of spatial information ^36^. The present results show that increases in pHPC-mPFC rsFC are beneficial to navigation task performance, thus contributing to the growing amount of literature emphasising the importance of information sharing between these two areas in spatial memory.

Post-glucose performance of the older participant was similar to that of the younger group. Conversely, in the younger group performance decreased slightly.

In young adults, the glucose facilitation effect is most readily observed under increased task difficulty ^21,37^ and divided attention ^38,39^. It is thus possible that the task demands in this study were not high enough to elicit an effect in the younger group. It is also the case that there are a number of studies where glucose modulation did not affect behavioural task performance ^2,40^, and there are isolated reports of decreased task performance after glucose ingestion ^18^. This may be a manifestation of complex interactions between glucose levels and task demands which make the effect more fragile in younger adults.

The reason for the difference between blood glucose levels 20minutes post-ingestion of the placebo drink are unknown, but they may reflect better glucoregulation as reflected intact insulin signalling after (false) nutritional load in the young group. Anticipatory hormonal response to flavouring (without glucose) have been observed before ^41^.

There were some limitations to this study. The sample sizes were relatively small, and future studies with larger sample sizes are needed in order to investigate the glucose facilitation effect especially with regard to mediating variables such as gender.

There is research suggesting that gender might be a contributing factor in the atrophy of the hippocampus ^42^ as well as a factor in the glucose facilitation effect ^43^. Future studies investigating the effects of gender are encouraged.

We used a hypothesis driven seed-based analysis to investigate the role of the anterior and posterior segments of the hippocampus in the glucose facilitation effect. This kind of analysis is influenced by the seed coordinates chosen. The seed coordinates in the present study were determined based on other investigations ^9^ to facilitate comparisons between studies. However other researchers have used different seed coordinates (e.g. ^44^) which may affect the results.

## Conclusion

This is the first study investigating functional connectivity of aHPC and pHPC after glucose load in a group of young and a group of elderly adults.

The results of the present study indicate the possibility that the pHPC is especially sensitive to pharmacological interventions, as we show that a simple glucose load modulates its connectivity and enhances cognitive performance in older adults.

Results also suggest that glucose modulated functional connectivity and cognitive performance more readily in older adults with impaired glucose regulatory ability. We further demonstrated the functional relevance of the changes in functional connectivity by relating gains in performance on a spatial memory task to increase in pHPC-mPFC connectivity.

## Supporting information

## Supplementary material accompanies this paper

### Contributions

A.S. and D.W. designed the experiment and supervised all aspects of the study. B.C. and R.P. contributed to the design of the study. R.P. collected and analysed the data, D.W. and B.C. contributed to data analysis. R.P. prepared the manuscript, D.W., A.S. and B.C. helped drafting and revising the manuscript. All authors have read and approved the manuscript.

## Acknowledgements

The study was partially supported by a scanning grant from Swinburne Neuroimaging (SNI) Facility, supported by the National Imaging Facility (NIF) under the National Collaborative Researcher Infrastructure Strategy (NCRIS).

## Competing interests

The authors declare no conflict of interest.

