## Supplementary material for "Functional connectivity of the anterior and posterior hippocampus: differential effects of glucose in younger and older adults"

### 1. Supplementary Procedures:

Detailed description of the two cognitive tests focusing on different domains of memory.

#### *1.1 Working memory*

Serial Sevens is a mental arithmetic task requiring the participant to repeatedly subtract seven from the previous number. Performance on this task has previously been found to be enhanced after glucose ingestion [1,2]

Instructions were displayed on a computer monitor, informing the participant to count backwards in sevens from a given starting number. Participants were instructed to ender responses using the numeric keyboard as quickly and accurately as possible. Participants were also instructed verbally that if they make a mistake they should carry on subtracting from the new incorrect number.

Participants were seated in front of a computer screen and a random number between 800 and 999 was presented as white numbers on a black background. The responses were made in the form of three digit numbers and each digit shown on the screen was replaced by an asterisk. After each response the participant had to press the enter key, which signaled the end of the response and cleared the three asterisks from the screen. In case of incorrect responses, subsequent responses were scored as correct if they were correct in relation to the new number.

The task duration was 2 minutes and performance was assessed using amount of correct subtractions.

#### *1.2 Spatial learning and memory*

Spatial learning performance was measured using a virtual analogue of the Morris Water Maze [3] which has been used extensively in rodents to study hippocampal-dependent spatial navigation. The virtual MWM has been shown to be sensitive to age and gender, as well as elicit hippocampal theta (4-8Hz) oscillatory activity that correlates with navigation performance. [4].

In each session, participants completed a set of 16 trials in which the escape platform was hidden (slightly submerged under the water) in the same location in relation to distal cues on the surrounding walls of the virtual environment. Starting locations at the pool's edge were pseudorandomly varied among the four cardinal points (N, S, E, W). The platform was made visible after 45 s if it had not been found. To test spatial memory, a probe trial was subsequently administered in which the platform was removed unbeknownst to participants. In this case, participants started from a novel position (e.g., NW) in order to assess the ability to generalize beyond immediate training experience. The platform was then moved to a new location, and participants completed an additional set of 12 hidden platform trials along with a second probe trial.

The dependent measure was heading error (square-root transformed) or angular deviation from a straight path to the platform's location on the probe trials. Due to computer malfunction during the placebo session, one participant's second probe trial data was not collected and thus replaced with the group-average heading error.

### 2. Supplementary Analysis

#### 2.1 Motion

Framewise displacement (maximum total and averages scan-to-scan) was calculated according to Power et al., (2012) [5] between-sessions and between-groups.

*Within-group* There was a within-group differences in average framewise displacement (FD) between glucose and placebo condition for the young group (placebo vs glucose: $t(1,13)=-2.602$; $p= 0.022$) but not for the older group (placebo vs glucose: $t(1,15)= -1.145$, $p= 0.27$). There were no within-group differences in maximum FD (young group placebo vs glucose: $t(1,13)= 0.432$, $p= 0.672$; elderly group placebo vs glucose: $t(1,15)= 0.449$, $p= 0.66$).

*Between groups* There was no between-group difference in maximum FD (young vs elderly on placebo: $t(1, 28)= -1.51$, $p= 0.142$; young vs elderly on glucose: $t(1,28)=-1.581$, $p= 0.131$). There were however differences between groups in average FD (young vs old on placebo: $t(1,28)= -2.48$, $p= 0.021$; young vs elderly on glucose: $t(1,28)= -2.181$, $p= 0.041$).

All group level analyses therefore included average framewise displacement as covariate at the second level.

#### 2.2 Calculation of partial grey matter volumes

To assess whether partial volume effects may influence the results, grey matter partial volumes for each of the ROIs were calculated. ROI specific GM volume was estimated from modulated, normalised grey matter volumes derived from unified segmentation of T1-weighted images as implemented in SPM12 following the procedure described by Pernet et al. 2009[6]. The same spatial normalisation parameters were used as for the preprocessing of rsfMRI data in spatial normalisation to MNI space.

Whole-brain voxel-wise rmANCOVA of resting state functional connectivity using left pHPC as a seed ROI were repeated with left pHPC partial volume as a covariate.

The cluster in the same area remained significant although smaller in size (MNI peak [+04 +24 +44]; Voxels (k)= 58) (**see Table S 2.2**).

**Table S 2.2**

**Whole brain voxel-wise rmANOVA of resting state functional connectivity with left posterior hippocampus controlling for pHPC volume**

| **Cluster *p*** | **Cluster size** | **Peak** | **MNI co-ord (peak)** | | | **Peak Region** |
|---|---|---|---|---|---|---|
| **(FWE-corr)** | ***k* voxels** | ***t*** | **x** | **y** | **z** | |
| .013 | 56 | -6.44 | 4 | 24 | 44 | Paracingulate gyrus R |

***Treatment x age-group interaction (young > older; Glucose > Placebo) for left posterior hippocampus (pHPC) seed (controlling for pHPC volume, two-sided contrast; 5000 permutations, FWE-corrected) to cluster in mPFC (encompassing anterior cingulate, paracingulate gyrus and superior frontal gyrus). The table displays cluster size p-value (FWE-corr), cluster size (k,) Peak t- value, Montreal Neurological Imaging (x, y, z) peak coordinates, peak region (Harvard-Oxford Atlas), R= right hemisphere.***
